# Focal Adhesion Kinase (FAK) inhibition induces membrane accumulation of aquaporin2 (AQP2) through endocytosis inhibition and actin depolymerization in renal epithelial cells

**DOI:** 10.1101/2024.10.08.617300

**Authors:** Asma Tchakal-Mesbahi, Jinzhao He, Shuai Zhu, Ming Huang, Kazuhiko Fukushima, Richard Bouley, Dennis Brown, Hua A. J. Lu

## Abstract

Cellular trafficking of the water channel aquaporin 2 (AQP2) is regulated by the actin cytoskeleton in collecting duct principal cells (PC) to maintain proper water balance in animals. Critical actin depolymerization/polymerization events are involved in both constitutive AQP2 recycling, and the pathway stimulated by vasopressin receptor signaling. Focal adhesion kinase (FAK) plays an important role in modulating the actin cytoskeleton through inhibiting small GTPases, and multiple studies have shown the involvement of FAK in insulin and cholesterol trafficking through actin regulation. To understand whether FAK contributes to water reabsorption by the kidney, we performed a series of in vitro experiments to examine the involvement of FAK and its signaling in mediating AQP2 trafficking in cultured renal epithelial cells. Our data showed that FAK inhibition by specific inhibitors caused membrane accumulation of AQP2 in AQP2expressing LLCPK1 cells by immunofluorescence staining. AQP2 membrane accumulation induced by FAK inhibition is associated with significantly reduced endocytosis of AQP2 via the clathrin-mediated endocytosis pathway. Moreover, AQP2 membrane accumulation induced by FAK inhibition also occurred in cells expressing the constitutive dephosphorylation mutant of AQP2, S256A. This was confirmed by immunoblotting using a specific antibody against phospho-serine 256 AQP2, supporting a phosphorylation independent mechanism. Finally, we demonstrated that inhibition of FAK caused reduced RhoA signaling and promoted F-actin depolymerization. In conclusion, our study identifies FAK signaling as a pathway that could provide a novel therapeutical avenue for AQP2 trafficking regulation in water balance disorders.

## Introduction

The water channel aquaporin2 (AQP2) is expressed in principal cells (PCs) of the collecting duct (CDs) and plays a crucial role in reabsorbing water along the CDs to produce concentrated urine (Fenton & Knepper, 2007), (Bouley et al., 2006), (Kanno et al., 1995), (Sasaki et al., 2000), (Sasaki, 2012). Significant advances have been made in understanding how AQP2 is regulated and the mechanisms governing its function in response to Vasopressin (VP). VP binds to its receptor, the vasopressin type 2 receptor (V2R), located on the basolateral membrane of CD principal cells, initiating a signaling cascade involving elevation of cyclic adenosine monophosphate (cAMP) and activation of protein kinase A (PKA). This series of events leads to the phosphorylation of AQP2 at serine-256, 264, and 269 and the dephosphorylation of serine-261, ultimately resulting in increased AQP2 accumulation on the apical membrane where it enables water reabsorption to occur. Phosphorylation of the AQP2 COOH-terminal serine 256 residue plays a dominant role in membrane accumulation of AQP2 compared to the other phosphorylated residues in the AQP2 c terminus (Katsura et al., 1997), (Fushimi et al., 1997), (Arthur et al., 2015), (Moeller et al., 2009).

Under non-stimulated conditions, AQP2 undergoes constitutive recycling between the plasma membrane and intracellular vesicles, independent of elevated intracellular cAMP levels. This cycling process occurs even in the absence of the antidiuretic hormone vasopressin (VP), and also when the S256 of AQP2 is substituted with alanine, an amino acid that cannot undergo phosphorylation (Nunes et al., 2008), (Lu et al., 2004). The importance of endocytosis in regulating this constitutive recycling pathway was first revealed by either depleting membrane cholesterol with methyl-β-cyclodextrin (mβCD) (Lu et al., 2004), (Russo et al., 2006) or by expressing a dominant-negative dynamin mutant (Sun et al., 2002). Both manipulations inhibit endocytosis without affecting exocytosis, and result in a rapid and substantial accumulation of AQP2 on the plasma membrane (Lu et al., 2004). Furthermore, we and others have shown that statins also cause AQP2 membrane accumulation by reducing its endocytosis via RhoA downregulation and actin depolymerization (Li et al., 2011), (Klussmann et al., 2001), (Tamma et al., 2001). However, this continuous equilibrium is disrupted by vasopressin, resulting in the accumulation of AQP2 on the membrane via increased exocytosis rates, inhibition of endocytosis, or a process involving both endocytosis and exocytosis. Similarly, AQP2 accumulation on the kidney principal cell’s plasma membrane in response to VP occurs through recycling between intracellular vesicles and the cell surface (Lu et al., 2004).

In vivo, VP modulates the levels of AQP2 present in the apical plasma membrane of kidney collecting duct cells. It does this partly by controlling the rates at which AQP2 undergoes endocytosis (internalization from the membrane) and exocytosis (insertion into the membrane). Regulation of endocytosis occurs by phosphorylating AQP2’s carboxyterminal phosphorylation site, primarily the serine residue at position 256, which reduces the rate of endocytosis (Brown, 2003). In addition, however, both the microtubule and actin cytoskeletons play crucial roles in AQP2 trafficking, and disruption of actin filaments alone results in AQP2 membrane accumulation by a combination of increased exocytosis and reduced endocytosis (Tamma et al., 2001), (Yui et al., 2012). This requires organized changes in cytoskeletal dynamics as well as the involvement of several associated proteins. Indeed, proteomic analyses have identified many cytoskeletal proteins and their regulatory interactors in both AQP2-bearing vesicles from inner medullary collecting duct (IMCD) and AQP2-containing exosomes isolated from urine (Pisitkun et al., 2004), (Barile et al., 2005).

Earlier studies have shown that RhoA inhibition is associated with the membrane translocation of AQP2 mediated by VP and cAMP. Moreover, AQP2 membrane accumulation can simply be induced by RhoA inactivation and actin depolymerization even in the absence of VP stimulation (Klussmann et al., 2001), (Valentt et al., 2005). Intriguingly, more recent studies have shown that integrins can signal from endocytic compartments and that FA proteins, including FAK, co-localize with endocytic proteins, which is required for integrin-dependent signaling (Alanko et al., 2015), (Nader et al., 2016), (Arriagada et al., 2019). The main function of the non-receptor protein tyrosine kinase, FAK, is to regulate vital cellular functions including proliferation, migration, differentiation, and apoptosis. In addition, it is known for its role in cellular processes related to cell adhesion, movement, or cell shape through regulating focal adhesion dynamics (Abedi & Zachary, 1995), (Schaller & Parsons, 1994). FAK is a convergent point in tyrosine phosphorylation pathways stimulated through the receptors for a variety of extracellular factors, including the regulatory peptides bombesin, vasopressin, endothelin, and angiotensin II, which act through G-protein-coupled seven transmembrane spanning domain receptors, and the regulatory lipids lysophosphatidic acid and sphingosine. Several of these factors also induce a rapid and striking reorganization of the actin filament network causing increased assembly of focal contacts and actin stress fiber formation (Abedi & Zachary, 1995).

Signaling through FAK mediates, the stabilization of microtubule interaction with the membrane, endocytosis, and vesicle trafficking including endocytosis of integrin complexes (Ezratty et al., 2005), (Chao et al., 2010), (Wang et al., 2011) and exocytosis of vesicles (Doucey et al., 2003), (Rosse et al., 2009),(Gupton and Gertler, 2010). In addition, FAK could be involved in the turnover of vesicles near the cleavage furrow (Almeida et al., 2000), (Guan, 2004), (Palazzo et al., 2004), (Avizienyte and Frame, 2005), (Wu et al., 2005). FAK-stimulated Src phosphorylation of endophilin inhibits dynaminmediated endocytosis (Wu et al., 2005), whereas FAK-stimulated Src phosphorylation activates endocytosis of cadherins (Avizienyte and Frame, 2005). Cai et al have shown that FAK is associated with protein trafficking such as during insulin secretion, where FAK deletion led to reduced insulin exocytosis likely by a defect in actin dynamics resulting in insufficient insulin granule trafficking (Cai et al., 2012). A recent study demonstrated an unsuspected role for FAK in controlling LDL-cholesterol delivery, specifically its late endosome trafficking and regulating the endosomal/lysosomal protein, Niemann–Pick C1 (NPC1) organelle dynamics through its coupling to the cholesterol transporter ORP2 (Takahashi et al., 2021). Zeigerer et al have shown that FAK-driven activation of Rab5 was associated with peripheral and enlarged early endosomes (Zeigerer et al., 2012). Importantly, it was previously reported that FAK activation and tyrosine phosphorylation are associated with RhoA activation and the formation of stress fibers (Schlaepfer & Mitra, 2004). In addition to affecting the activity of Ras, Rac, and Rho, FAK can influence the function of Cdc42 through binding and phosphorylation of the Cdc42 effector Wiskott–Aldrich syndrome protein N-WASP (neuronal WASP), which leads to actin cytoskeleton regulation through activation of the ARP2/3 complex (Wu et al., 2004), (Ridley et al., 2003). Our laboratory recently revealed a crucial role of the Arp2/3 complex in AQP2 exocytosis (Liu et al., 2021). Finally, FAK along with paxillin and talin, regulate intracellular cytoskeleton dynamics (Mitra et al., 2005). Building on these findings, we set out to elucidate a possible role of FAK in regulating AQP2 transport, and we report here that FAK modifies AQP2 translocation through actin cytoskeleton regulation.

## Materials and Methods

### Cell culture

LLCPK1 cells stably expressing wild type c-myc-tagged aquaporin-2 (LLCPK1-AQP2 cells) (Katsura et al., 1995), LLCPK1-AQP2 cells stably expressing secreted soluble yellow fluorescent protein (LLCPK1-AQP2-ssYFP cells) (Nunes et al., 2008), LLCPK1AQP2 cells stably expressing alanine 256 mutant c-myc-tagged aquaporin-2 (LLCPK1AQP2-S256A) (Nunes et al., 2008), (Rice et al., 2012) were all cultured at 37°C with a 5% CO_2_ atmosphere in DMEM + 10% FBS + 2 mg/ml glucose. Cells were grown either on cover slips (Electron Microscopy Sciences, Hatfield, PA) or on standard P6 polystyrene Falcon culture dishes (Corning Science Mexico) to 70 - 80% confluence. (Chemicals are listed in Supplemental Table1).

### Endocytosis and exocytosis assay

For the endocytosis assay, LLC-AQP2 cells were grown on coverslips placed in a 12-well cell culture plate to 70-80% confluence (Corning Science Mexico). After incubation in serum-free DMEM for 1 h, LVP, VS-4718 (a FAK inhibitor) and DMSO as a control were added and incubated with cells for 30 minutes. Cells were then washed in ice-cold PBS twice, and then incubated with tetramethylrhodamine transferrin at a final concentration of 20-25 mg/ml in 1% BSA / no phenol red DMEM for 20-30 minutes. Cells were washed 3 times immediately with ice-cold PBS, fixed with 4% paraformaldehyde (PFA) in PBS and visualized by fluorescence microscopy (Nikon 80i). For the exocytosis assay, LLCAQP2-ssYFP cells were cultured and treated as above, and then exocytosis of ssYFP secreted into the culture medium was measured by quantifying the increase in fluorescence in aliquots of culture medium. The specific protocol was previously described in detail (Nunes et al., 2008).

### Immunofluorescence staining and immunoblotting

Cells were treated as described above, then fixed in 4% PFA in PBS for 20 min. Fixed coverslips were permeabilized in 0.02% Triton X-100/PBS for 4 min, washed three times with PBS, and blocked in 1% BSA/PBS for 30 min. The coverslips were incubated with primary antibodies for 1 h at room temperature or overnight at 4°C. After washing with PBS three times, coverslips were incubated in secondary antibody for 45 min at room temperature, and after 3 more washes with PBS the coverslips were mounted with Vectashield solution (mounting medium with DAPI, Vector Laboratories Inc., Burlingame, CA). Evaluation was done using a Nikon 80i or Bio-rad confocal microscope. Image J (NIH; http://rsb.info.nih.gov/ij/) software was used for image analysis. Background corrections and contrast/brightness enhancement were performed identically for all images in the same experiment. For immunofluorescence, AQP2 was detected using undiluted culture medium containing mouse anti-c-myc IgG produced by the hybridoma cells (MYC 19E10.2 [9E10] ATCC® CRL1729TM) followed by a secondary anti–mouse IgG conjugated to Alexa-488 (10 *μ*g/ml; Invitrogen). For cell lysate preparation, cells were grown on P6 plates and treated with different chemicals as described above, then rinsed 3 times with ice-cold PBS and lysed with a cell scraper in 150 μl lysate buffer containing 8 mM NaF, 0.5% NP-40, 4 mM Na3V04, 0.1% Triton X-100, 0.03% NaN3 and 1% protease inhibitor cocktail from Biotool.com (Houston, TX). Samples were incubated on ice, passed more than 5 times through a 27G×1/2 syringe and then centrifuged at 12,000 rpm at 4°C for 10 min to remove cell debris. The protein concentration in the lysate was measured with the Pierce® BCA Protein Assay Kit from Pierce Biotechnology (Rockford, IL). After adding 3X lithium dodecyl sulfate (LDS) sample buffer (Life Technologies, Carlsbad, CA) the samples were used for western blot as described before (Rice et al., 2012). AQP2 was detected using a goat polyclonal antibody, sc-9882 (1:500 dilution) from Santa Cruz Biotechnology (Dallas, TX), mouse anti-GAPDH monoclonal antibody AM4300 (at 1:10,000) from Ambion/Thermo Fisher Scientific (Lu et al., 2004). (Antibodies are listed in Supplemental Table 2).

### Cold block assay experiment

LLC-AQP2 cells were grown on 12×12mm glass coverslips to 70%-80% confluence. In this cold block assay, cells were treated with different chemicals and placed in 20°C water bath to block the recycling of protein/vesicles, that causes the appearance of perinuclear patch structures. Cycloheximide (10 μg/ml) was added to the culture medium 60 min before experiments to inhibit newly synthesized proteins. Coverslips were taken out at different time points, fixed with 4% PFA and immunofluorescence staining was performed as previously described (Lu et al., 2004).

### RhoA activity Assay

LLCPK1-AQP2 cells were cultured in standard P10 polystyrene Falcon culture dishes (Corning Science, Mexico) to 70-80% confluence, washed with ice-cold PBS, and harvested in lysis buffer with a cell scraper. Equivalent protein amounts of lysates (300 g800 g) were then incubated with 50 g Rhotekin-RBD beads (Cytoskeleton, Inc) at 4°C on a rotator for 1 hour. The beads were pellet by centrifugation at 5000g then washed by removing the supernatant. Then, beads were suspended in 2x Laemmli sample buffer and boiled for 2 minutes before being analyzed with SDS-PAGE and Western blot using anti-RhoA antibodies.

### Intracellular cAMP measurement

Cells plated on 96 well plates were treated with LVP and VS-4718 as above, then analyzed using an ELISA based cAMP DetectX Direct Cyclic AMP ELISA Kit (Arbor Assays) according to the manufacturer’s protocol. LLCPK1-AQP2 cells were treated with 50 mM indomethacin overnight before exposure to both treatments (Deen et al., 1997). Absorbance values at 450 nm were measured using the DTX880 plate reader. The results were adjusted by cell number and analyzed by a two-tailed Student’s t-test.

### F-actin depolymerization assay

8×10^4^ cells were plated on 24 well plates and incubated in normal DMEM for 2 days (LLCPK_1_). Cells were treated with 50 µM indomethacin overnight prior to use (Deen et al., 1997). DMEM was replaced with HBSS and cells were incubated for 2 hours. After treatment with VP/FK or latrunculin A as above for various times, HBSS buffer was removed and cells were incubated in binding buffer containing phalloidin (20 mM KH_2_PO_4_, 10 mM PIPES, 5 mM EGTA, 2 mM MgCl_2,_ 4% PFA, 0.1% TritonX-100, 250 nM rhodamine phalloidin) for 15 min at room temperature. Negative controls to estimate background autofluorescence were prepared using binding buffer lacking rhodamine-phalloidin. Cells treated with binding buffer were washed 4 times in PBS, incubated in 300 µl (in each well) methanol overnight at −20°C to extract bound rhodamine-phalloidin. The extracted rhodamine-phalloidin fluorescence was read using a Beckman DTX-880 multi-plate reader (excitation 535 nm, emission 595 nm). F-actin content values were expressed as relative fluorescence after subtraction of the negative-control values. Each fluorescence value was expressed as relative fluorescence unit (RFU)/well and data were analyzed using a two-tailed Student’s *t*-test (Yui et al., 2012).

### Statistics

Statistics were performed with Prism software (GraphPad, La Jolla CA). Data for each treatment were first compared for significance with a one-way ANOVA. Differences in means were then compared between control and treatment at each time point using the student’s T-test (two tailed). Each experiment was repeated at least three times. Statistical significance was determined at a p value <0.05.

## Results

### Inhibiting FAK signaling caused membrane accumulation of AQP2 in cultured cells

FAK inhibition with VS-4718 caused membrane accumulation of AQP2 in cultured cells. We first examined the effect of VS-4718 on AQP2 trafficking in cultured LLC-PK1 cells stably expressing AQP2 (LLC-AQP2). Treatment with VS-4718 led to a significant accumulation of AQP2 in the plasma membrane domain while AQP2 was distributed in a relatively diffuse vesicular pattern in control cells (Fig. 1A). In cells treated with LVP, membrane accumulation of AQP2 was clearly detected. VS-4718-induced membrane accumulation of AQP2 with 1µM VS-4718 occurred after 30 min of treatment (Fig. 1). Importantly, there was no significant cytotoxicity observed in treated cells, which is consistent with other reports (de Jonge et al., 2019), (Mohanty et al., 2020).

**Fig. 1.**
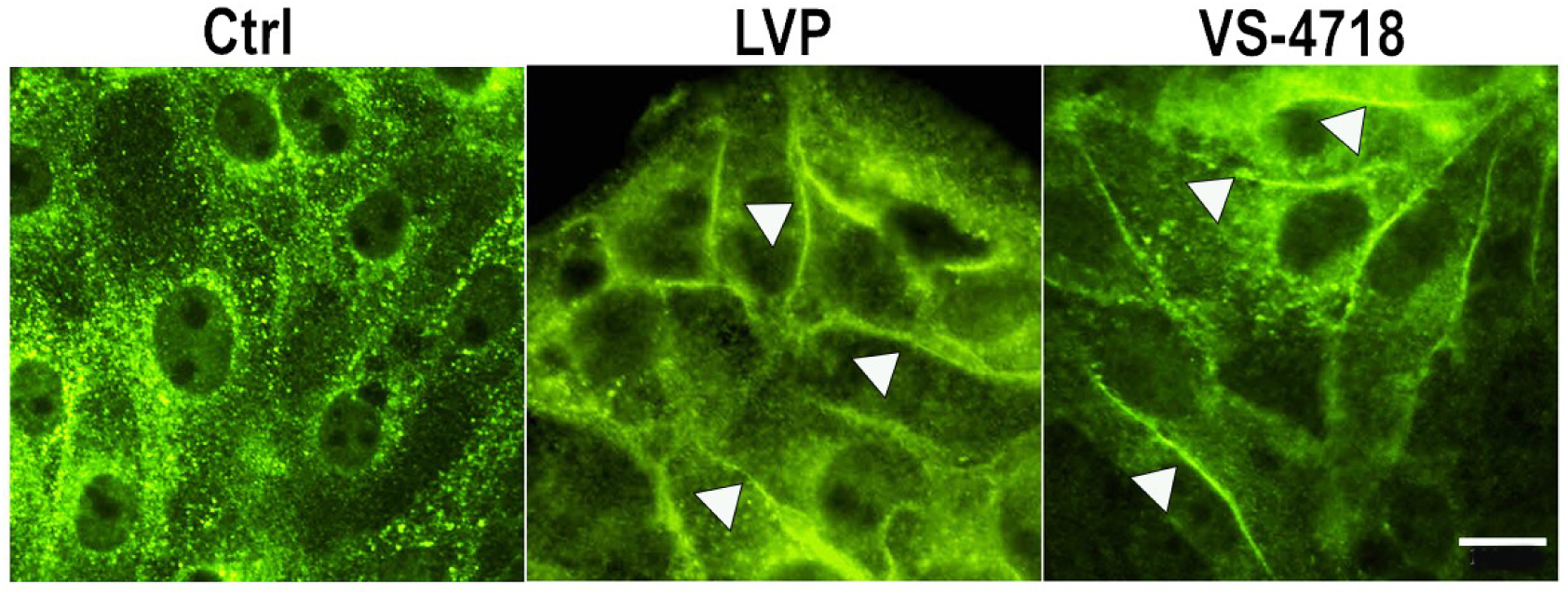
A. FAK inhibition-induced membrane accumulation of AQP2 in LLCPK1-AQP2 cells. Immunofluorescence staining of AQP2 with anti-c-Myc antibody in LLCPK1AQP2 cells treated with 1µM VS-4718 for 30 min indicates a strong membrane accumulation of AQP2 as shown by LVP treatment (10 nM, 15 min).

### FAK inhibition caused reduced endocytosis of AQP2 in cells

We examined whether increased membrane accumulation of AQP2 by VS-4718 is due to altered endocytosis. AQP2 is known to be endocytosed via the clathrin-mediated pathway (Brown, 2003). We performed a rhodamine-conjugated transferrin endocytosis assay. VS4718 treatment blocked the endocytosis of rhodamine-transferrin and led to increased membrane accumulation of rhodamine-transferrin signal compared with the control condition (Fig 2A). As expected, MbCD also blocked endocytosis significantly (Fig. 2B).

**Fig. 2.**
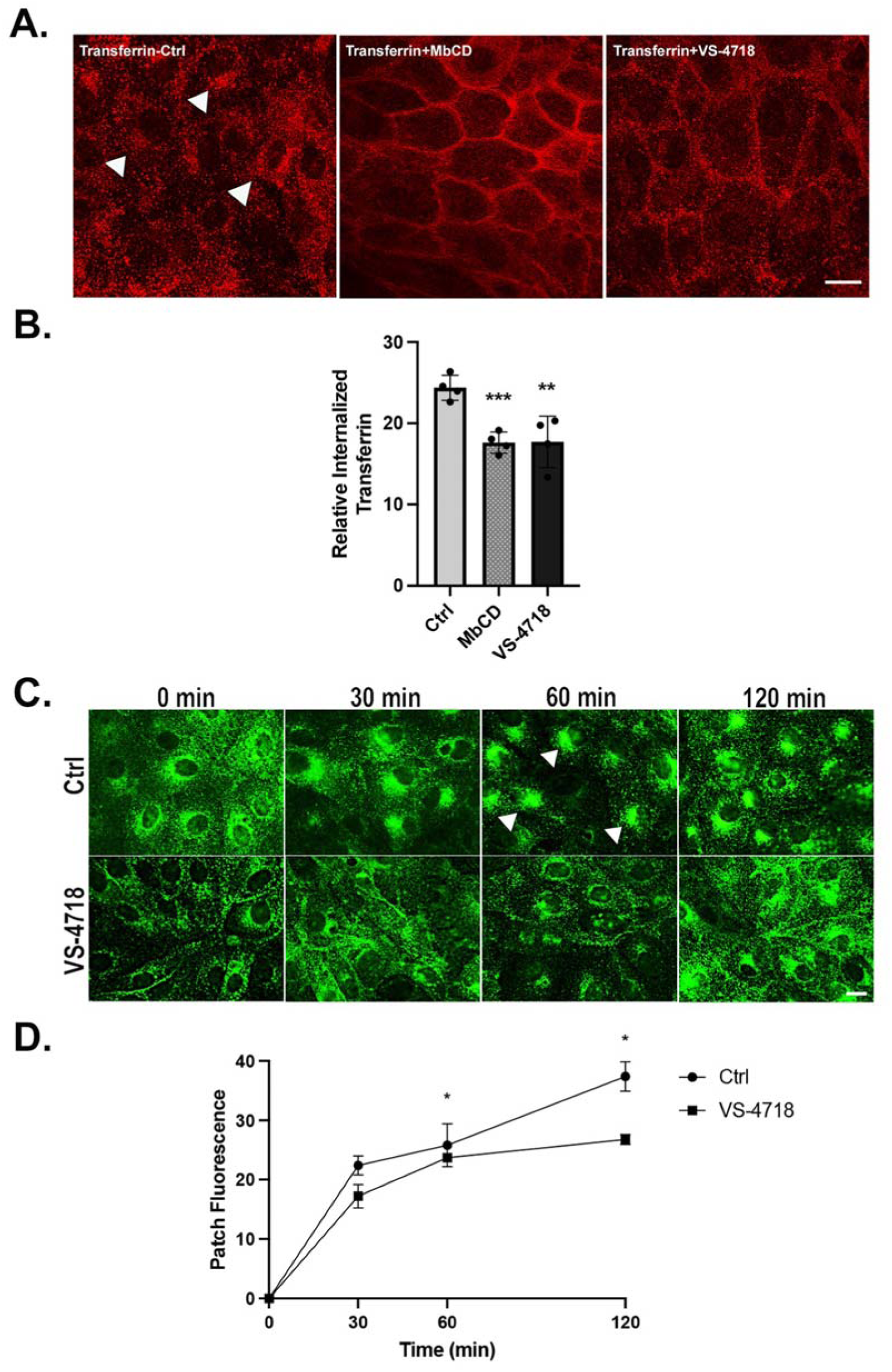
VS-4718 reduces AQP2 endocytosis. **A.** Clathrin-mediated endocytosis was followed using rhodamine-conjugated transferrin on LLCPK1-AQP2 cells. VS-4718 blocked rhodamine-transferrin endocytosis and led to increased membrane accumulation. Methyl-b-cyclodextrin (MbCD) is a positive control to block endocytosis. **B.** Relative internalized transferrin quantification by measuring mean pixel intensity of patch using ImageJ software. Student’s t-test was performed to examine significance of treated groups vs. control (**P < 0.01; ***P < 0.001). **C.** Dynamic intracellular accumulation of AQP2 in “cold block” experiment. VS-4718 treatment inhibited the usual perinuclear accumulation (arrows in Ctrl) of AQP2 after cold block at 20°C. Scale bar = 10 µm. **D**. Fluorescence signal intensity of intracellularly accumulated AQP2 after cold block experiment in LLCPK1-AQP2 cells shows a significant delay in patch accumulation after treatment with VS-4718. Student’s t-test was performed to examine significance of treated groups vs. control (*P < 0.05).

Following this, we specifically examined the endocytosis of AQP2 in LLCPK1-AQP2 cells using a cold block assay that is routinely used to measure the rate of AQP2 internalization (Rice et al., 2012), (Arthur et al., 2015). Incubating cells at 20°C inhibits the exit of proteins from the trans-Golgi network and their subsequent transport to the cell plasma membrane (Griffiths et al., 1985). The internalized proteins accumulate in the perinuclear region, forming a so-called “perinuclear patch” (Arthur et al., 2015). In the absence of protein synthesis, the size and density of the perinuclear patch reveal the rate of internalization of the respective membrane protein. In control cells, an AQP2 perinuclear patch was clearly observed ∼60 min after cold block, and the patch signal plateaued ∼120 min. VS4718 treatment led to delayed endocytosis of AQP2; therefore, the formation of an AQP2 peri-nuclear patch was significantly delayed compared with the control, indicating an inhibited endocytosis process (Fig.2C). The fluorescence intensity of AQP2 perinuclear patches over time was quantified and analyzed (Fig. 2D). It showed that the decrease of the AQP2 perinuclear patch in VS-4718-treated cells was significantly delayed suggesting that VS-4718 impedes AQP2 endocytosis/internalization in cells.

### FAK inhibition induced an increase in AQP2 exocytosis

To explain further the AQP2 accumulation on the membrane when FAK is inhibited, we performed the exocytosis assay using a soluble secreted yellow fluorescent protein (ssYFP) as a surrogate marker of AQP2 exocytosis in LLC-AQP2 cells (Nunes et al., 2008). We found that both LVP and VS-4718 significantly increased exocytosis of ssYFP by 20% compared to control (Ctrl) (Fig. 3).

**Fig. 3.**
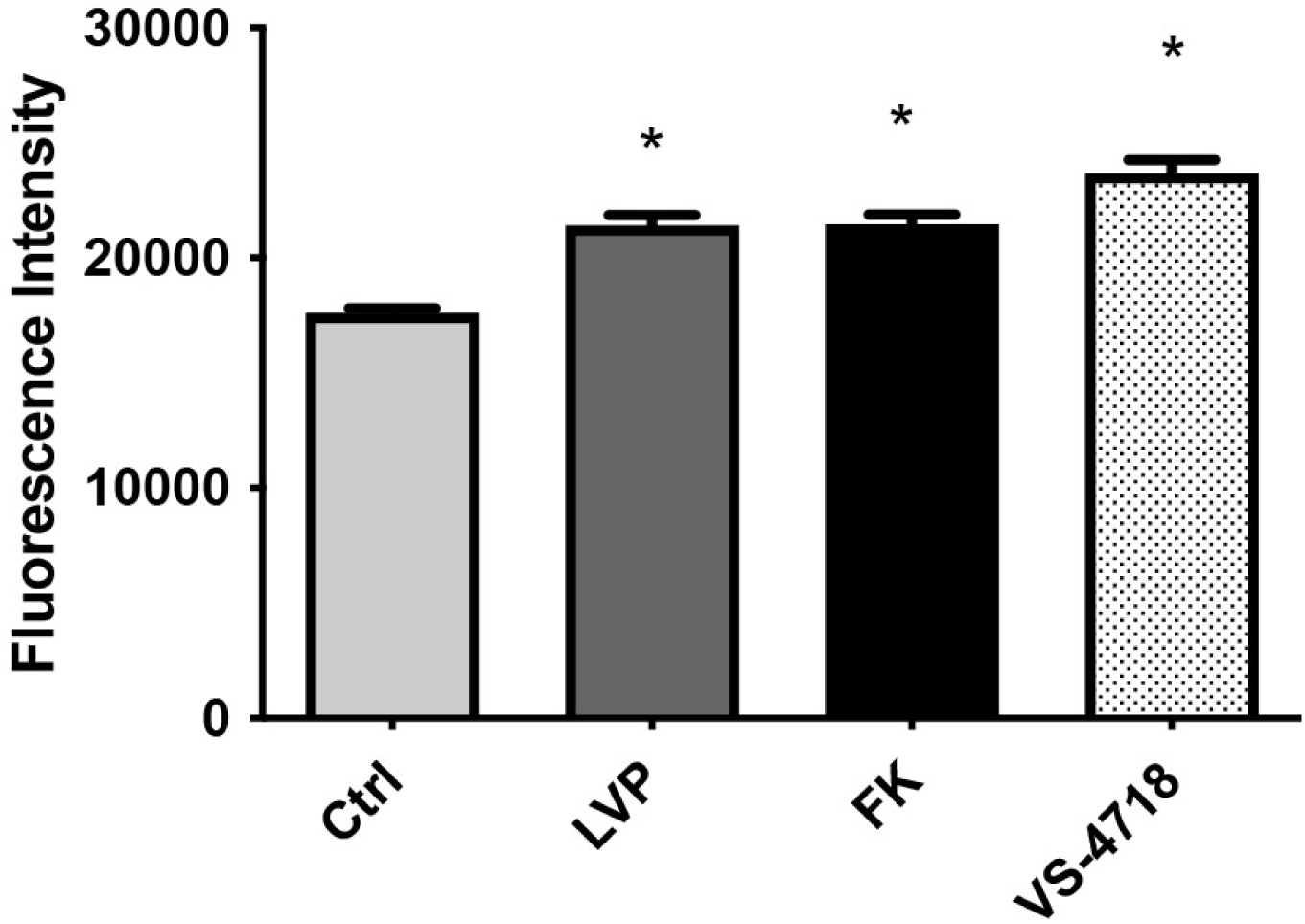
VS-4718 increases AQP2 exocytosis. Relative fluorescence of ssYFP in LLCPK1-AQP2-ssYFP cells as quantified. All experiments were repeated at least three times. Ctrl, control. VS-4718 treatment induces a significant increase in ssYFP intensity indicating an increase in AQP2 exocytosis rate. Bar values represent means + SE. Student’s t-test was performed to examine significance of treated groups vs. control (*P < 0.05), n ≥ 3).

### Membrane accumulation of AQP2 induced by FAK inhibition occurs independently of S256 AQP2 phosphorylation and cAMP signaling

It is well known that AQP2 trafficking in response to vasopressin requires its phosphorylation at its COOH-terminal residues, particularly, phosphorylation of AQP2 at its serine-256 residue, a critical for its trafficking and regulation (Lu et al., 2008), (Brown et al., 2008), (Nielsen et al., 1995), (Moeller et al., 2009), (Fenton and Knepper, 2007). To confirm that FAK inhibition induces AQP2 accumulation at the membrane in an S256 phosphorylation-independent way, we examined the effect of FAK inhibition on AQP2 trafficking in LLCPK1-256A cells, a cell line expressing a mutation of AQP2 at its S256 phosphorylation site by the substitution of the amino-acid serine to alanine. Methyl-βcyclodextrin (MβCD) treatment blocked the endocytosis of AQP2 and led to its membrane accumulation compared with the control. Similarly, VS-4718 caused significant membrane accumulation of AQP2 in LLCPK1-256A cells confirming the trafficking of AQP2 without S256 phosphorylation (Fig. 4A). Our continued investigation using specific phosphorylated antibody against serine 256 AQP2 clearly shows that in contrast to LVP, which induces S256 phosphorylation, VS-4718 treatment of LLC-AQP2 cells did not result in S256 phosphorylation (Fig. 4B) These findings suggest that VS4718-induced AQP2 membrane accumulation is unlikely to involve AQP2 phosphorylation.

**Fig. 4.**
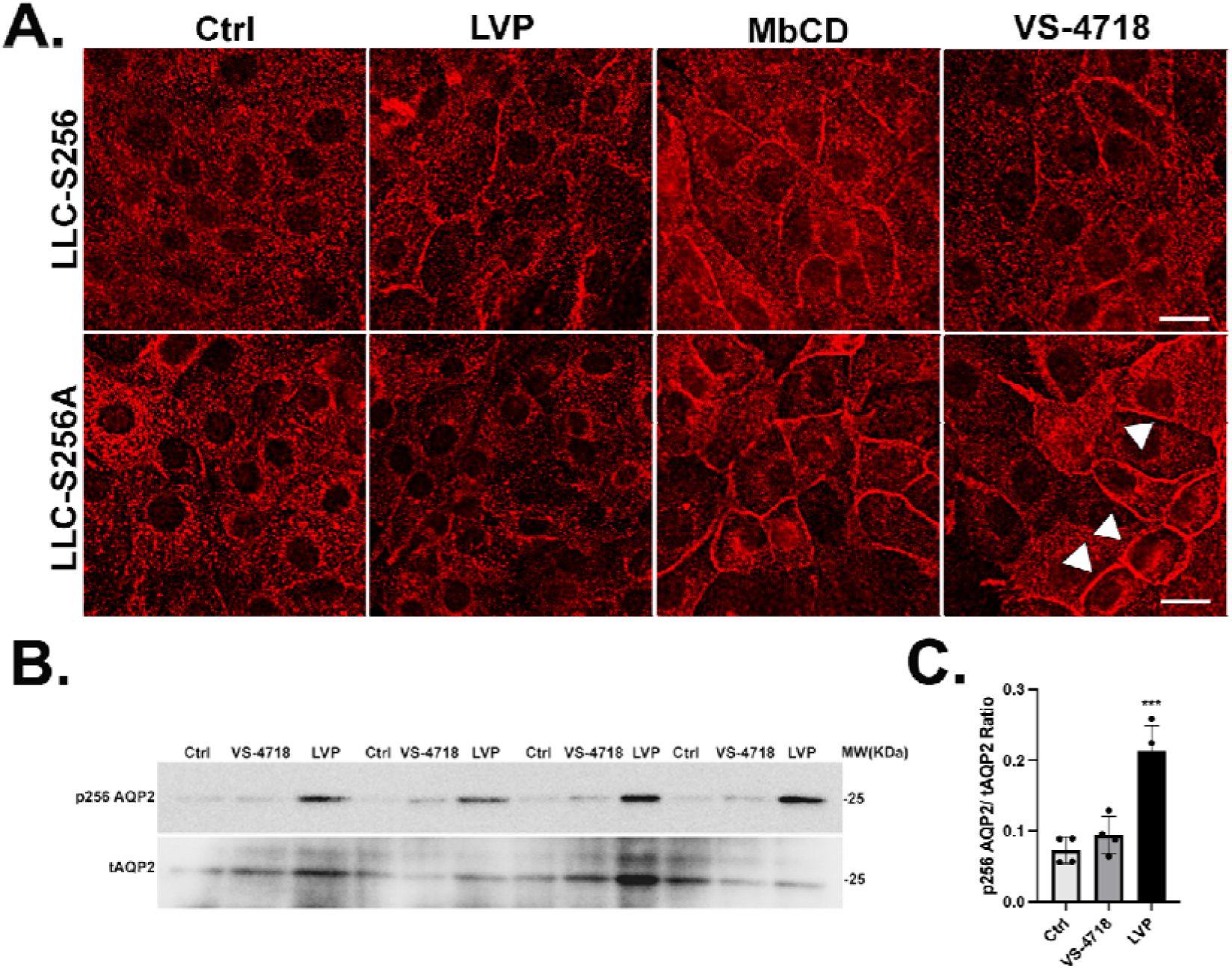
VS-4718 induces AQP2 membrane accumulation without phosphorylation at serine 256. **A.** Membrane accumulation of AQP is seen in cells expressing either WTAQP2 or the S256A phosphorylation mutant. Methyl-beta-cyclodextrin (MbCD) is a positive control to block endocytosis, scale bar=10 µm. **B.** Western Blot of tAQP2 and pS256 indicates that VS-4718 does not induce AQP2 trafficking through the canonical, VP-dependent pathway. After detecting p256AQP2 with specific anti p256AQP2 antibody, membranes were stripped and reblotted with AQP2 antibody **C**. Bar graph represents ratio of p256 AQP2 and total AQP2 band intensities of 4 independent experiments (mean ± SE, n=4, *** is p<0.001).

### FAK inhibition does not increase intracellular cAMP in LLCPK1-AQP2 cells

To investigate further if VS-4718 involves the cAMP/PKA pathway to shift AQP2 to the membrane in LLCPK1 cells, we tested whether FAK inhibition induced an increase in intracellular cAMP concentration in LLC-PK1 cell lines. In LLCPK1-AQP2 cells. VS-4718 did not induce an increase in intracellular cAMP whereas LVP increased cAMP as expected (Fig. 5A). In addition, western blotting showed no increase of cAMP-response element binding protein (CREB) phosphorylation after FAK inhibition in LLCPK1-AQP2 cells unlike, vasopressin which produced a strong signal compared to non-treated cells (Fig. 5B).

**Fig. 5.**
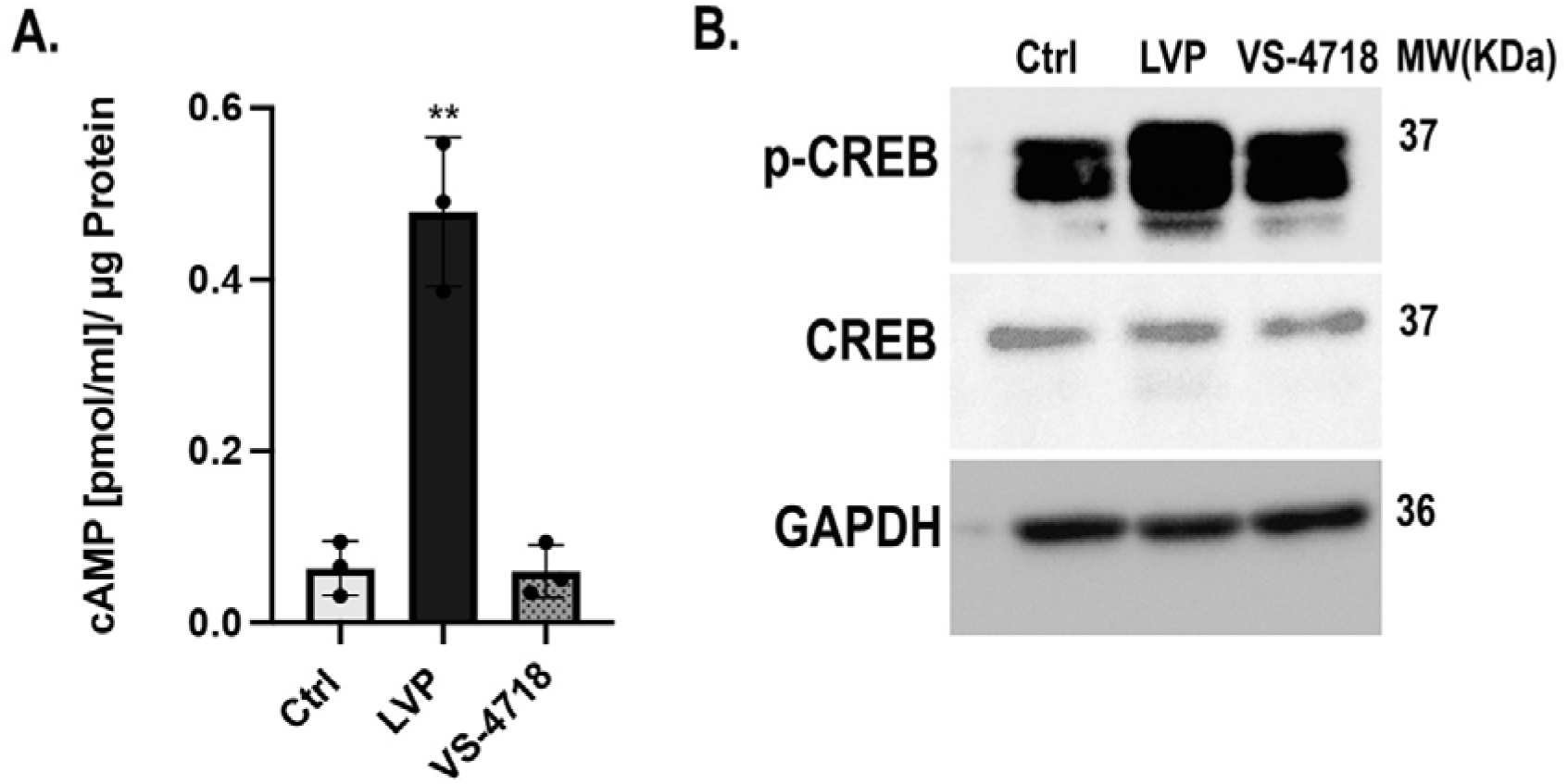
VS-4718 induces AQP2 membrane accumulation without increasing cAMP and CREB phosphorylation in LLCPK1 cells. **A.** Intracellular cAMP concentration measurement in LLCPK1-AQP2 cells after treatment with LVP and VS-4718, with ELISA. **B**. Western Blot Analysis of phosphorylated CREB indicates that FAK inhibition does not induce an increase of phosphorylated CREB signal compared to vasopressin treatment in LLCPK1-AQP2 cells.

### FAK inhibition induced AQP2 membrane accumulation by modifying the actin cytoskeletal network and inhibiting RhoA

Actin polymerization and depolymerization are recognized as crucial factors in regulating the dynamic trafficking of AQP2. We performed F-actin quantification assays using LLCPK1-AQP2 cells. Cells were treated with VP/FK (LVP 20 nM, FK 10 mM) and 1uM of VS4718 for the indicated times, then subjected to F-actin quantification. In LLC-PK1-AQP2 cells, the F-actin content was significantly decreased by about 10-15% after VP/FK treatment compared to untreated cells. In a similar way, VS-4718 induced an F-actin depolymerization of the same magnitude in LLC-PK1 AQP2 cells compared to controls. Latrunculin A, used as a positive control resulted in a 50% decrease in F-actin concentration (Fig. 6A). these data are similar to the previously reported effect of VP on F-actin depolymerization in LLC-PK1 cells that express AQP2 (Yui et al., 2012).

**Fig. 6.**
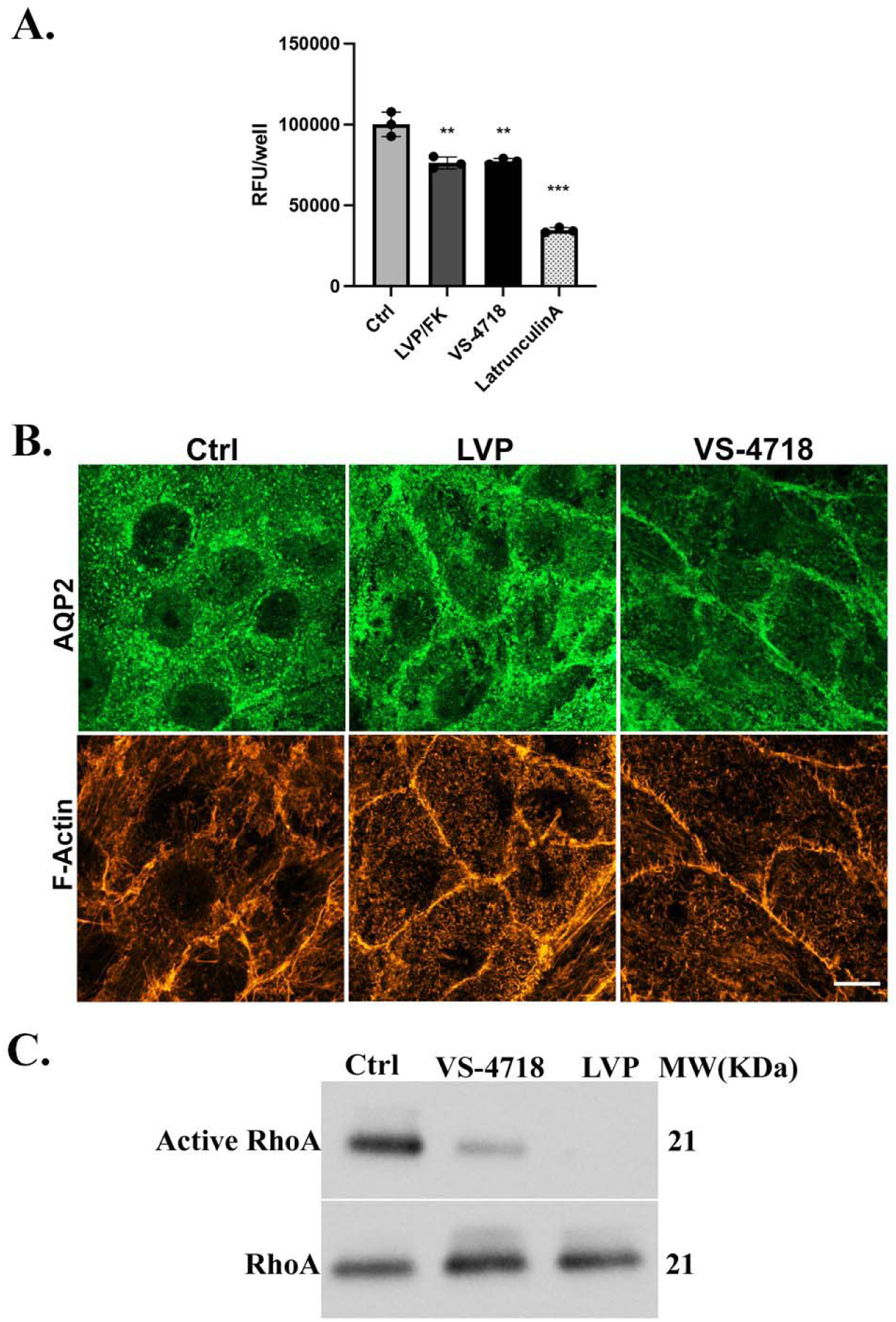
VS-4718 induces AQP2 membrane accumulation by modulating F-actin and inhibiting RhoA. **A.** F-actin quantification assay using rhodamine-phalloidin binding shows that VS-4718 and LVP cause a significant 20% decrease in F-actin compared to controls. where Latrunculin A, an actin depolymerizing agent, induces a decrease of 50% F-actin-used as a positive control here. **B**. Immunofluorescence using phalloidin to detect F-actin and c-myc to detect tagged AQP2, after treatment with vasopressin and VS-4718. VS-4718 and LVP caused AQP2 membrane accumulation, and some decrease in peripheral F-actin, scale bar=10 µm. **C**. RhoA pulldown assay using GST-RBD beads that bind to active RhoA in LLCPK1-AQP2 cells shows that VS-4718 reduces Active RhoA in a similar way to vasopressin.

We then conducted immunofluorescence staining of F-actin in cells exposed to LVP or the FAK inhibitor, VS-4718, utilizing rhodamine-conjugated phalloidin. The administration of LVP or VS-4718 both led to the accumulation of AQP2 in the cell membrane but at this level of resolution, major changes in the F-actin distribution were not readily detectable. More detailed study will be required using higher resolution imaging to examine this in the future (Fig. 6B). However, compared to untreated cells, the amount of GTP-RhoA was strongly reduced after FAK inhibition, consistent with an attenuation of Rho activity (Fig. 6C). As expected, a drastic reduction in GTP-bound cellular RhoA was observed in LLCPK1-AQP2 cells treated with vasopressin, known to inactivate protein RhoA.

## Discussion

Our major new finding here is that FAK inhibition causes membrane accumulation of AQP2 in LLC-PK1 renal epithelial cells. Membrane accumulation of AQP2 by FAK inhibition is independent of AQP2 phosphorylation at serine 256 and does not require the elevation of intracellular cAMP. This membrane accumulation of AQP2 induced by FAK inhibition involves decreased actin depolymerization and inactivation of the small GTPase RhoA. Membrane accumulation could be due to an increase in AQP2 exocytosis and/or a decrease in its endocytosis, leading to membrane accumulation. Using our Rhodamine-transferrin endocytosis assay (Lu et al., 2004), we confirmed that AQP2 is accumulated in the membrane after FAK inhibition and the rate of endocytosis was decreased in these cells, indicating that AQP2 internalization is reduced by FAK inhibition. The cold block assay also showed that FAK inhibition induces a significant delay in the perinuclear accumulation of AQP2 internalization, which is associated with reduced endocytosis as demonstrated by the Rhodamine-transferrin assay. However, an increased rate of exocytosis was also revealed after FAK inhibition with VS-4718 using our secreted yellow fluorescent protein (ssYFP) assay, in a similar way as seen in LVP-response (Brown, 2003), (Brown et al., 2008), (Nunes et al., 2008).

Although AQP2 S256 phosphorylation through the cAMP/PKA pathway is required to cause apical plasma membrane accumulation of AQP2 in kidney principal cells in response to VP (Brown, 2000), (Deen, 2001), (Nielsen et al., 2002), (Sasaki et al., 2000), (Bouley et al., 2000), it is evident now that this translocation can also be induced by other maneuvers that appear to be independent of PKA (Lu et al., 2004), (Russo et., 2006). Our results now reveal that AQP2 membrane accumulation induced by VS-4718 does also does not require the VP/cAMP pathway since FAK inhibition induced membrane accumulation of AQP2 in the mutant cell line LLCPK1-256A and also does not trigger the phosphorylation of AQP2 at S256, a crucial site for AQP2 trafficking and regulation. The phosphorylation status of AQP2 at serine-256 in the presence of FAK inhibition was monitored using a phosphorylation-specific antibody.

Regarding the mechanism by which FAK inhibition affects AQP2 trafficking, we have shown here that FAK inhibition regulates actin dynamics by reducing the activity of the small GTPase RhoA. The actin cytoskeleton plays an important role in regulating AQP2 trafficking through exocytosis and endocytosis (Klussmann et al., 2001) and actin depolymerization alone can lead to the cell-surface accumulation of AQP2 in the absence of any hormonal stimulus (Klussmann et al., 2001), (Valenti et al., 2005), (Tamma et al., 2003), (Mamuya et al., 2016). Transition between actin polymerization and depolymerization is a master key of AQP2 trafficking which is regulated by RhoA (Mamuya et al., 2016), (Noda et al., 2004), (Sasaki et al., 2014). RhoA is widely involved in vesicle and protein trafficking by reorganizing stress fibers (Noda et al., 2004), (Pochynyuk et al., 2007), (Stirling et al., 2009). Moreover, RhoA inhibition and actin depolymerization are associated with VP- and cAMP-mediated membrane translocation of AQP2. In addition, AQP2 membrane accumulation can simply be induced by RhoA inactivation and actin depolymerization even in the absence of VP stimulation (Tamma et al., 2001), (Tamma et al., 2003). Our current data support these earlier observations and now implicate FAK in this process.

Importantly, the membrane accumulation of AQP2 after FAK inhibition appears to be mainly a result of the inhibition of AQP2 endocytosis as was shown by the cold block assay. Although phosphorylation of AQP2 is required for VP-induced membrane accumulation of AQP2, VS-4718-mediated membrane accumulation of AQP2 seems to require a different mechanism independently of AQP2 phosphorylation at its residue S256. We demonstrated that RhoA is an important player in AQP2 transport upon FAK inhibition. Indeed, FAK inhibition induces AQP2 membrane accumulation through RhoA inhibition in a similar manner to vasopressin but via a pathway other than cAMP/PKA. In summary, our study has uncovered an important and potent effect of FAK on regulating AQP2 trafficking in cells bypassing the VP-V2R (cAMP/PKA) signaling pathway and orchestrating changes in dynamic cytoskeletal architecture.

## Acknowledgements

This work was supported by the National Institutes of Health (NIH) grant DK096915 (H. J.L.) DK096586-1 (D.B.) and diversity supplement DK096586-10A1S1(to A.M.). Additional support for the Program in Membrane Biology Microscopy Core comes from the Boston Area Diabetes and Endocrinology Research Center (DK057521) and the Massachusetts General Hospital (MGH) Center for the Study of Inflammatory Bowel Disease (DK043351).

## Notes

### Competing Interest Statement

The authors have declared no competing interest.

