## Supplemental Table 1, Supplemental Table 2 for "Focal Adhesion Kinase (FAK) inhibition induces membrane accumulation of aquaporin2 (AQP2) through endocytosis inhibition and actin depolymerization in renal epithelial cells"

**Supplemental Table 1: List of chemicals**

| Chemical | Company |
|---|---|
| Lysine-Vasopressin | Sigma |
| VS-4718 (HY-13917/PND-1186) | MedChem Express |
| GSK | MedChem Express |
| DH | MedChem Express |
| Latrunculin A | CalBiochem |
| Methyl-β-cyclodextrin | Sigma Aldrich |
| Forskolin | Biotechne Tocris |
| Alexa Fluor® 555 Phalloidin | Cell Signaling Technology |
| Tetramethylrhodamine transferrin | Life Technologies (Carlsbad, CA) |

**Supplemental Table 2. List of antibodies**

| Antigen | Manufacturer | Catalog | References | Dilution (WB) | Dilution (IF/IHC) |
|---|---|---|---|---|---|
| RhoA | Cell Signaling | 2117T | PMID: 37730597 | 1:1000 | |
| GAPDH | Applied Biosystems | 4300 | PMID: 32434945 | 1:10000 | |
| p-CREB | Cell Signaling | 9198S | PMID: 32223896 31822662 | 1:1000 | 1:400 |
| CREB | Cell Signaling | 9197S | PMID: 32223896 | 1:1000 | |
| P256 AQP2 | Affinity | 3673 | PMID: 15509592 | 1:1000 | |
| P269 AQP2 | Cell Signaling | 3674 | PMID: 15509592 | 1:1000 | |
| P261 AQP2 | Rockland antibodies | 612-401 D08 | PMID: 7508187 | 1:1000 | |
| AQP2 | Santa Cruz | 9882 | PMID: 22218592 | 1:1000 | |
| $Na^+/K^+$ ATPase | Abcam | 76020 | PMID: 35235339 | 1:1000 | |
